# Diversity of Mammal Fauna in a Remnant of the Atlantic Forest in the Middle of a Megalopolis

**DOI:** 10.64898/2026.06.05.730359

**Authors:** Mariana Malzoni Furtado, Aspen Caim Fernandes Gonçalves, Melissa de Sá Fernandes, Grasiela Edith Oliveira Porfírio, Maristela Martins de Camargo

## Abstract

A remnant Atlantic Forest fragment within São Paulo’s urban environment supports six mammal species, with Didelphis being most abundant, while maintaining natural temporal activity patterns and successful reproduction, demonstrating the conservation value of small protected areas within megacities for preserving native wildlife despite intense anthropic pressure. The Forest Reserve of the Institute of Biosciences (FRIB) is a remnant of the Atlantic Forest located at the University of São Paulo, city of São Paulo *campus*, in Brazil. Despite urban pressures such as altered food resources, pollution, pathogens, and interactions with invasive and domestic species, several wildlife species persist, demonstrating adaptability to urban environments. This study aimed to sample the mammal species diversity of FRIB using camera traps. A total of 367 records of six mammal species were obtained over 107 days of sampling, between June and November 2024, resulting in 7,872 hours of sampling effort. The species recorded were *Callithrix jacchus, Callithrix penicillata, Didelphis aurita, Didelphis albiventris*, and unidentified species of the orders Chiroptera and Rodentia. *Didelphis sp*. was the most recorded genera with 201 records. These results highlight the significance of this forest fragment in protecting wild animals in an environment facing intense anthropic pressure.

**Resumo:** A Reserva Florestal do Instituto de Biociências (FRIB) é um remanescente da Mata Atlântica situado na Universidade de São Paulo, em São Paulo, Brasil. Apesar das pressões urbanas, como mudanças nos recursos alimentares, poluição, patógenos e interações com espécies invasoras ou domésticas, diversas espécies de animais selvagens continuam existindo, evidenciando sua capacidade de adaptação aos ambientes urbanos. O objetivo deste estudo foi amostrar a diversidade de espécies de mamíferos da FRIB através de armadilhas fotográficas. Durante um período de 107 dias de amostragem, de junho e novembro de 2024, foram obtidos 367 registros de seis espécies diferentes de mamíferos, totalizando 7.872 horas de esforço. As espécies registradas foram *Callithrix jacchus, Callithrix penicillata, Didelphis aurita, Didelphis albiventris*, além de espécies não identificadas das ordens *Chiroptera* e *Rodentia. Didelphis sp*. foi o gênero com maior número de registros (n=201). Esses resultados ressaltam a importância desse fragmento florestal na proteção da fauna selvagem em um ambiente que enfrenta intensa pressão antrópica.

## Introduction

The Forest Reserve of the Institute of Biosciences (hereafter referred to as FRIB), also known as the “Matinha da Biologia”, is a remaining area of the Atlantic Forest within the *campus* of the University of São Paulo. Despite its location in the urban center of São Paulo, the reserve provides a favorable environment for various native wildlife species.

The presence of wild animals in urban environments is becoming increasingly frequent worldwide. While some species are unable to survive, others exhibit great adaptive flexibility and can thrive in anthropized environments, even with reduced natural areas (Magle et al., 2019). However, urban areas also offer stressful perturbations, such as changes in food resources, exposure to pathogens, acoustic and visual stress, air pollutants, and interactions with new species including invasive or domestic animals (Soulsbury and White, 2015; Appel and Porfirio, 2020; Perry et al., 2020; Rucco et al., 2020).

Studies conducted worldwide on various *taxa* have demonstrated that urbanization leads to the loss of species with specialized diets, breeding sites, or specific habitat requirements, while generalist species tend to adapt more successfully (Soulsbury and White, 2015). Additionally, behavioral flexibility, such as a high tolerance for disturbances, appears to be a crucial trait for thriving in urban environments (Lowry et al., 2012).

According to the book “Fauna and Flora of the *Campus*”, published in 2017 by Jane Kraus and collaborators, the University of São Paulo harbors a rich biological diversity, including black-eared opossums (*Didelphis aurita*), small rodents, lizards, thrushes, marmosets (*Callithrix* sp.) and sloths *(Bradypus* sp.), among others. Recent observations on iNaturalist (www.inaturalist.org), a citizen science platform that shares biodiversity information, have reported sightings of black-eared opossums, marmosets, green-billed toucans (*Ramphastos dicolorus*), yellow-headed woodpeckers (*Celeus flavescens*) and more. However, no academic studies or detailed works about the species living in the reserve were found.

Camera traps provide an efficient, non-invasive approach for the continuous sampling of wildlife, enabling the investigation of ecological and conservation questions (Lyra-Jorge et al. 2008; Trolliet et al. 2014). In addition, there is a positive outlook surrounding next-generation camera traps, as they allow the formulation of previously infeasible questions encompassing a broader range of species and a greater variety of environments (Delisle et al. 2021). As part of a research project by the Institute of Biomedical Sciences of the University of São Paulo to evaluate and adapt a non-invasive biosurveillance tool for collecting saliva from marmosets, opossums and wild rodents in natural environments (Camargo et al., 2026), a survey of the fauna of the FRBI using camera traps was conducted. Therefore, we aimed to present the results of the fauna survey through the analysis of species richness and composition obtained from camera trapping data.

## Material and Methods

The main campus of the University of São Paulo (USP) comprises an area of 791.97 hectares, with 124 hectares or 15.66% of its territory occupied by ecological reserves. The Forest Reserve of the Institute of Biosciences (FRIB) covers an area of 10.2 hectares with a 30 meters slope. This fragment is a remnant of the Atlantic Forest and has the Tejo stream at its headwaters. The area does not allow public access, only authorized staff can conduct research and extension projects within the area.

Between June 26 and November 6, 2024, four digital camera traps (Bushnell HD) were installed in the FRIB. Three sampling campaigns were conducted: 1) Ground Sampling 1 from June 26 to July 15; 2) Ground Sampling 2 from July 29 to August 12; and 3) Arboreal Sampling 1 and 2 from August 27 to November 6 (Figure 1).

**Figure 1.**
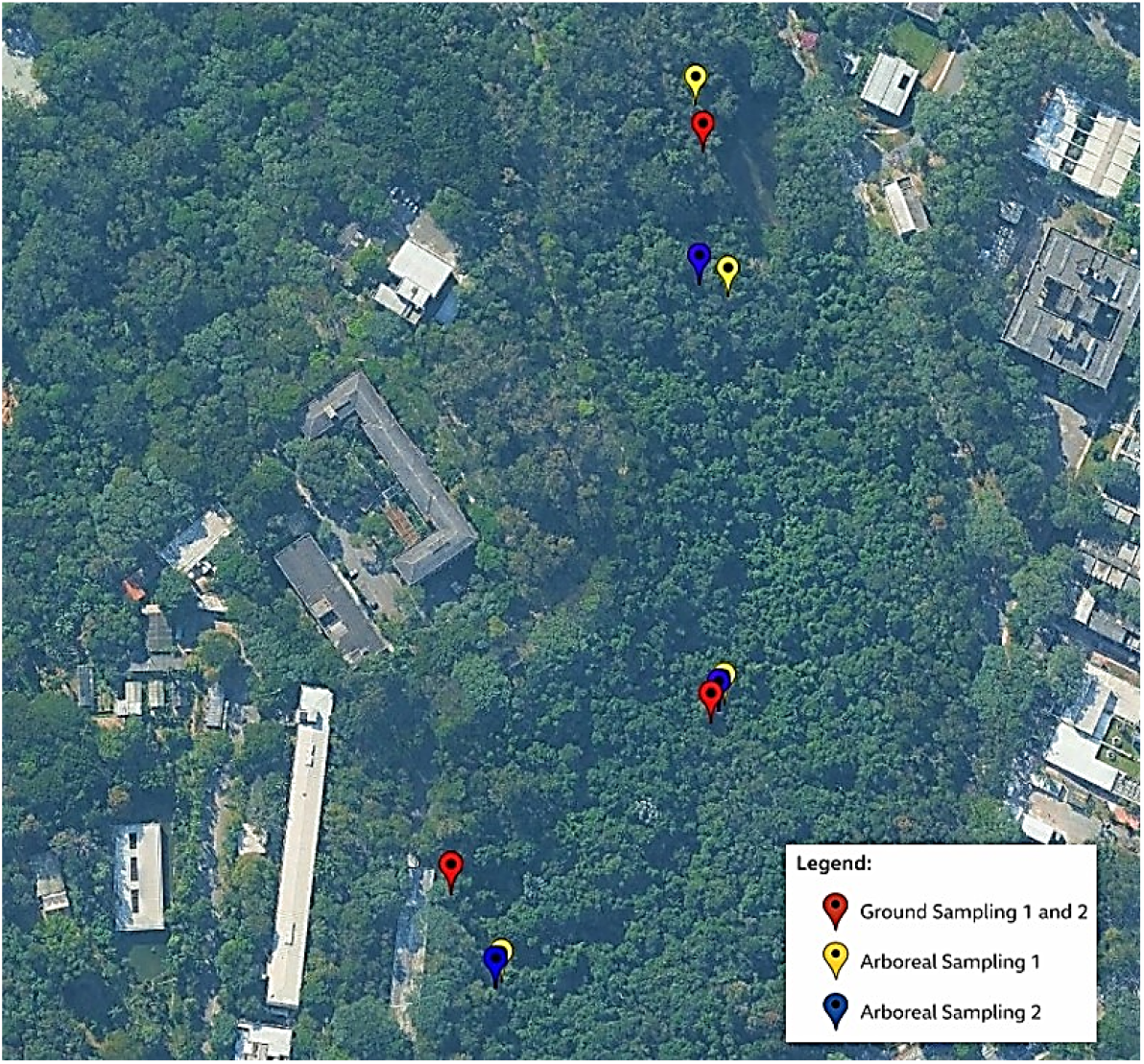
Location of the camera traps installed in the Forest Reserve at the Institute of Biosciences of the University of São Paulo.

**Figure 2.**
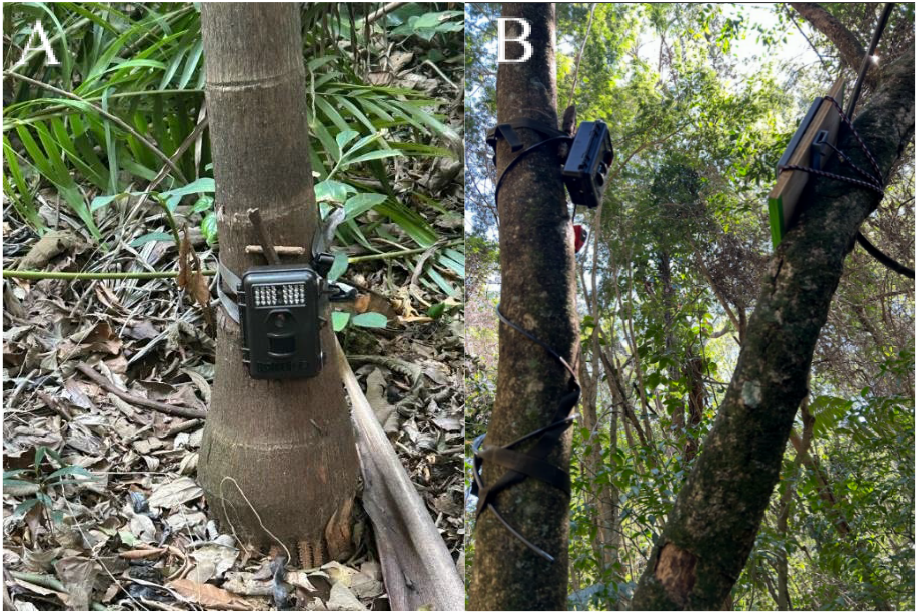
Camera traps installed to record (A) terrestrial and (B) arboreal fauna in the Forest Reserve at the Institute of Biosciences of the University of São Paulo.

The camera traps were positioned in trees or close to the ground near existing trails. During the first two campaigns, three to four camera traps were set up at the same spots at an average height of 20 to 40 centimeters from the ground, to capture images of terrestrial fauna. In the third campaign, three camera traps were placed at the canopies of trees, approximately 1.70 to 2.30 meters from the ground, to capture images of arboreal fauna. The location of the cameras changed throughout this campaign, resulting in Arboreal Sampling 1 and Arboreal Sampling 2. The camera traps had infrared sensors activated by motion. Cameras were checked periodically (daily in sampling 2, weekly in samplings 1 and 3) to replace batteries, memory cards and perform necessary maintenance. These camera traps were installed as part of a research project to document wildlife interaction with saliva collectors that have attractive odors (Camargo et al., 2026; https://sites.usp.br/swab/). Therefore, some of the animals recorded might not exhibit their natural behavior, as they were attracted by the presence of the attractive odors.

For the first sampling campaign, the camera traps were programmed to record three photos and one 30 second video. In samplings 2 and 3, the cameras were programmed to record one 15 second video. The capture effort was calculated in hours by multiplying the number of camera traps by the number of hours sampled (24 hours/day) (Srbek-Araujo & Chiarello, 2007).

Photo and video records were analyzed considering their independence. Records of the same species obtained after a five-minute interval were considered independent. Consecutive records of the same species, captured in the same trap within less than five minutes, were excluded from the analysis, and this set of consecutive images was computed as a single independent record (Srbek-Araujo & Chiarello, 2007). Whenever possible, individuals were identified to the species level based on recordings. However, due to image quality of videos/photos, or records showing only parts of the animal’s body, some individuals were identified by their Order, Family or Genus, while others remained unidentified.

Morphological features visible in the camera trap images were used for species identification. To distinguish between *Didelphis* species, the color of the inner ear was a defining characteristic: black for *D. aurita* and white for *D. albiventris*. For *Callithrix* species, the color of the ear tuft was used for identification, white for *Callithrix jacchus* and black for *Callithrix penicillata*. Other characteristics like length and weight were not considered in this study.

## Results and Discussion

A sampling effort of 107 days (or 7,872 hours) was conducted, resulting in a total of 393 independent records. Out of these, 341 were from the Mammalia Class (consisting of 8 species) and 49 were from the Aves Class (comprising 9 species). There were three instances where the animals could not be identified (Tables 1 and 2, Figure 3).

**Table 1.**
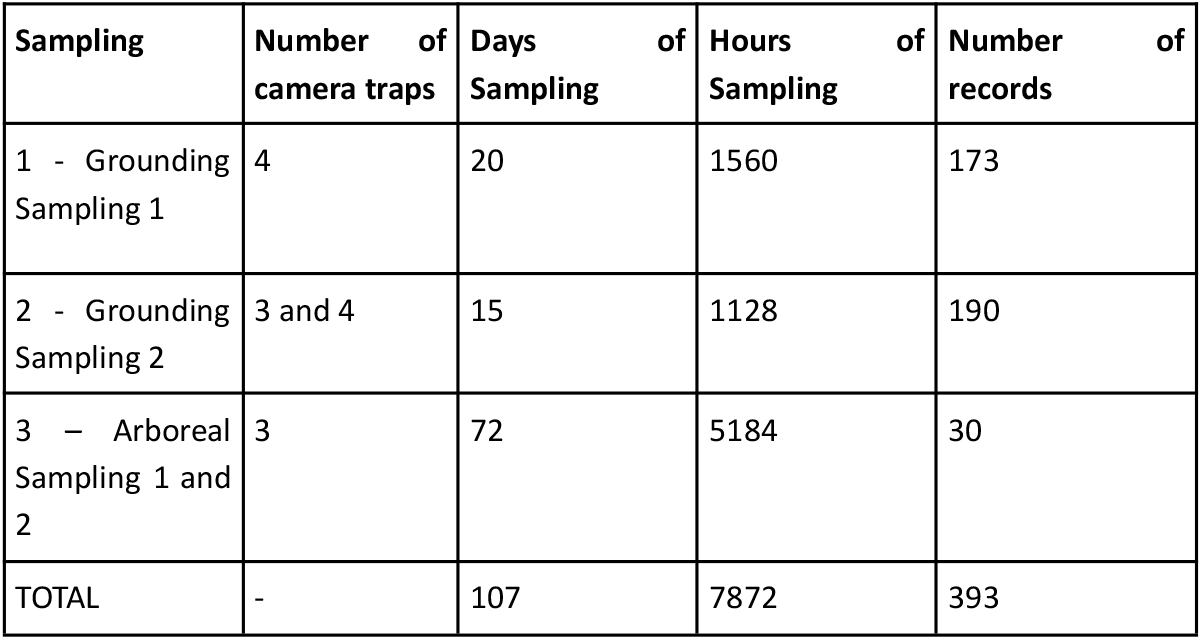
Sampling effort with camera traps in the Forest Reserve of the Institute of Biosciences of USP from June to November 2024.

**Table 2.** Species of mammals and birds recorded in camera traps installed in the Forest Reserve of the Institute of Biosciences of USP between June and November 2024.

| Class recorded | Species recorded | Conservation Status* | Sampling |  |  | TOTAL |
| --- | --- | --- | --- | --- | --- | --- |
|  |  |  | Terrestrial 1 | Terrestrial 2 | Arboreal 1, 2 |  |
| Mammalia | *Didelphis* spp. | - | 100 | 95 | 6 | 201 |
|  | *Didelphis aurita* | LC | 42 | 81 | 2 | 125 |
|  | *Didelphis albiventris* | LC | 6 | 8 | 1 | 15 |
|  | Chiroptera | - | 0 | 0 | 5 | 5 |
|  | Rodentia | - | 4 | 0 | 1 | 5 |
|  | *Callithrix* spp. | - | 0 | 0 | 5 | 5 |
|  | *Callithrix jacchus* | LC | 0 | 0 | 1 | 1 |
|  | *Callithrix penicillata* | LC | 0 | 0 | 10 | 10 |
|  | **TOTAL** |  | **152** | **184** | **31** | **367** |
| Aves | Ave | - | 17 | 3 | 1 | 21 |
|  | *Aramides saracura* | LC | 5 | 0 | 0 | 5 |
|  | *Troglodytes musculus* | LC | 1 | 0 | 0 | 1 |
|  | *Turdus rufiventris* | LC | 1 | 5 | 0 | 6 |
|  | *Mimus saturninus* | LC | 2 | 0 | 0 | 2 |
|  | *Penelope obscura* | LC | 2 | 0 | 0 | 2 |
|  | *Furnarius rufus* | LC | 0 | 1 | 0 | 1 |
|  | *Mesembrinibis cayennensis* | LC | 2 | 1 | 0 | 3 |
|  | *Columbina talpacoti* | LC | 3 | 5 | 0 | 8 |
|  | **TOTAL** |  | **33** | **15** | **1** | **49** |
| Unidentified |  |  | 1 | 2 | 0 | 3 |

\*Conservation status from Brazilian Environmental Agency. LC= Least Concern

Species richness estimates that mix species- and higher-rank detections can be misleading; we therefore report them separately to avoid inflating species counts. Among mammals, the following species were recorded: *Didelphis aurita*, an endemic species of the Atlantic rainforest (Assis, 2011) (n=125); *Didelphis albiventris*, typically found in the Cerrado and Caatinga (Nascimento, 2015) (n=15); *Callithrix jacchus* (n=1); and *Callithrix penicillata* (n=10). In some cases, it was not possible to identify the species, and some individuals were identified by their genus (n=201 *Didelphis* spp., n=5 *Callithrix* spp.). Bats and wild rodents were identified only by their order. We used encounter rates (independent records per 100 camera-hours) to compare detections across campaigns with unequal effort and configurations. Event counts may be biased by camera height, settings, and the use of olfactory attractants; therefore, we interpret results as detections/encounter rates rather than true abundance. The most recorded species in terrestrial sampling was *Didelphis aurita*, and in arboreal sampling, it was *Callithrix penicillata* (Table 2). Among birds, 8 species were identified, and 21 recorded birds were not identified (Table 2). For rodents, visible morphological characteristics, such as long vibrissae and a tail ending in a long tuft of hairs point to the genus *Rhipidomys* (climbing mice). This genus includes two species native to the Atlantic Forest: *Rhipidomys tribei* and *R. itoan* (Costa, et al. 2011). Currently, *R. tribei* is classified as Endangered, while *R. itoan* is classified as Least Concern. However, it was not possible to identify the rodents to the species level based on the records obtained. The conservation *status* shown in the table was obtained from ICMBio’s SALVE platform (www.salve.icmbio.gov.br).

The records of *Didelphis aurita*, accounted for 36% of the total sampling, and can be attributed to the increase in individuals in fragmented areas where there are few or no predators (Assis, 2011). This is also true for the FRBI, as native predators are no longer found in the area. On the other hand, *Didelphis albiventris*, is predominantly found in the Cerrado, and thus, a small number of records in the Atlantic Forest was expected.

*Callithrix jacchus* and *C. penicillata* can easily adapt to urban areas. They exhibit a great ability to occupy these habitats due to their generalist diet and high reproductive rate (Elvas et al. 2023). In São Paulo city, marmosets (*Callithrix* sp.) have been frequently found and their population has been increasing in one of the largest metropolis in the world (Fioratti, 2023). *Callithrix jacchus* is considered an invasive species in São Paulo. Originally native to the Northeast it was introduced into the Southeast region by trafficking and illegal housing as pets. This introduction has led to environmental imbalances, including competition for resources and hybridization with *Callithrix penicillata* (Conti, 2023).

The diversity of species in FRIB was higher in the central region of the reserve, particularly in areas influenced by the Tejo stream, compared to the more open and disturbed areas near the entrance gates and the lake. A greater diversity of mammals was also observed in arboreal sampling. Although we spotted a larger number of bird species during fieldwork, not all species were captured on camera traps.

During October and November, camera traps recorded *Didelphis* sp. joeys, suggesting that the end of winter (July) and the beginning of spring (September) coincide with the species’ breeding season (Monteiro-Filho et al, 2006). On three separate occasions, individuals were recorded walking in pairs, suggesting a breeding season in July/August.

Opossums have crepuscular or nocturnal habits (Monteiro-Filho et al, 2006), as confirmed by our study. The species’ activity was concentrated between 4:30 pm and 6:30 am. In contrast, marmosets are diurnal animals (Auricchio, 1995) and were active in the RFIB during the day, between 10 am and 3:45 pm. Although olfactory baits were used, we cannot rule out effects on detection timing; thus, activity windows should be interpreted cautiously. We did not estimate survival, recruitment, body condition, or disease; thus, activity alignment and juvenile detections alone do not demonstrate successful adaptation.

In urban centers like São Paulo, it is increasingly common to observe wild animals adapting to urbanized areas. Notable examples include marmosets, opossums, capybaras and different species of birds (Augusto, 2023; Fioratti, 2023). These animals often scavenge for food in trash cans, gardens or parks, which encourages their proximity to inhabited areas. However, the growing population of these animals in urban spaces brings significant challenges. Cohabiting humans unfamiliar with these species, inadequately feed them, which can increase their reproduction rates, consequently increasing their presence in urban areas (Hemétrio, 2011, Takahata and Kutsukake, 2025). On the other hand, their increased presence might lead to human-animal conflicts (Soulsbury et al., 2015). Additionally, wild animals are exposed to road accidents, pathogen transmission by humans and domestic animals, and behavioral changes caused by constant human proximity (Nantes et al., 2021; Barreto et al., 2024; Santo et al., 2025). In this context, protected areas can be a good solution. They provide a safe refuge allowing animals to reproduce and maintain natural behaviors.

## Conclusion

The results of this camera trap survey indicate that FRIB reserve functions as a refuge where recorded mammals exhibit activity windows consistent with the literature; however, we did not infer successful adaptation or population viability due to lack of demographic and health data.

## Acknowledgments

This study was funded by Wild Animal Initiative (USA)(SG-23-030) and Scholarships Programs PUB (University of São Paulo). We thank the RFIB management committee, especially Paulo Diaz, and Dr. Gabriel Marroig for their valuable comments, and Dr. Renata Pardini for making trap cameras available for this study.

## Notes

### Competing Interest Statement

The authors have declared no competing interest.

